# Changing seasonality and phenology mediate the effects of migration across meta-ecosystems

**DOI:** 10.64898/2025.12.03.692017

**Authors:** Lara Chaouat, Florian Altermatt, Tianna Peller

## Abstract

Migration is a ubiquitous process that links ecosystems with distinct seasonal dynamics, transferring biomass and species interactions across space. Despite being widely altered by global change, studies commonly overlook the interaction of seasonal characteristics and bidirectional migration on species coexistence and biomass production across meta-ecosystems. We developed a mathematical model to study how migration interacts with key characteristics of seasonality—amplitude of variation and length of summer—to influence migrant persistence, consumer coexistence, and biomass production. Our findings demonstrate that seasonal characteristics mediate the effect of migration on coexistence and biomass production across meta-ecosystems. However, the effects strongly depend on migration timing: phenological mismatches can reduce biomass at local and meta-ecosystem scales and lead to the extinction of migratory and non-migratory consumers. Our study highlights how migration and seasonality interact to shape community structure and ecosystem function across scales, emphasizing the importance of system-level approaches for studying ecological outcomes of global change.

## Introduction

Migration is a ubiquitous phenomenon in nature, with billions of animals moving between ecosystems each year (Bauer and Hoye, 2014; Milner-Gulland et al., 2011). Ungulates, for instance, track seasonal shifts in vegetation (Harris et al., 2009), zooplankton perform daily vertical migrations to balance predation risk and foraging (Bandara et al., 2021), and many waterfowl species undertake perilous long-distance journeys (Newton, 2007). Decades of research have shown that migration benefits migratory species by providing better foraging opportunities, protection from predators, or greater reproductive success (Shaw, 2016). Migration, however, also serves as an ecological link connecting the biodiversity and functions of otherwise disconnected ecosystems (Bauer and Hoye, 2014; Gounand et al., 2018; Subalusky et al., 2017)–a role that remains relatively understudied (Bauer and Hoye, 2014; Hansen et al., 2020; Peller et al., 2023).

A substantial body of research has established that migration is a strategy to contend with periodic changes in ecosystem conditions, in particular, seasonality (Milner-Gulland et al., 2011). Seasonality is characterized by a variety of factors, including changes in temperature, precipitation, and photoperiod, which can influence the availability of food or breeding sites, and the abundance of competitors and predators (McMeans et al., 2015). As ecosystems can differ in their seasonal characteristics, such as the intensity, timing, and length of seasons (Conover, 1992), organisms can migrate from one ecosystem to another in search of more favorable conditions. Although the influence of seasonal characteristics on local ecosystems is increasingly recognized (Doležal et al., 2019; Hernández-Carrasco et al., 2025; McMeans et al., 2015; White and Hastings, 2020), the interaction between seasonal characteristics and migration, and its implications for the dynamics, diversity, and functioning of ecosystems across scales, remains largely unexplored (Hutchison et al., 2020).

Global change is altering seasonality and the ecological processes centered around it (Ernakovich et al., 2014; Hernández-Carrasco et al., 2025; Walther et al., 2002). In particular, evidence suggests global change is intensifying seasonal variations (Cohen et al., 2021) and modifying the duration of seasons (Lin and Wang, 2022), with projections suggesting longer warm periods and shorter cold periods. Such shifts in seasonality can affect migratory species, who often depend on seasonal cues to prompt their migration between ecosystems (Berthold, 1996; Doiron et al., 2015; Post and Forchhammer, 2007). This can lead to phenological mismatches where the timing of migration does not coincide with key ecological events (Bairlein, 2016; Cohen et al., 2018), such as peaks in resource availability (Beard et al., 2019). For example, in some avian species, migration is triggered by photoperiod at wintering ecosystems, making their arrival dates at breeding or nesting ecosystems often inflexible. This can lead to a mismatch between the arrival dates of migrants at breeding sites and the peak of insect abundance, which are a vital resource for migrants (Both and Visser, 2001). As migration mismatches are emerging around the world (e.g.; Clausen and Clausen (2013); Mayor et al. (2017); Post and Forchhammer (2007)), it raises crucial questions about how migratory species shape biodiversity and ecosystem functions and how these roles are being altered with global change. Moreover, while some species are not fast enough to adapt to shifting seasonal cues, others adjust their migratory behavior accordingly (e.g.; Haest et al. (2020); Harris et al. (2009); Lameris et al. (2018); Ward et al. (2016)), raising important questions about how such changes in migration will interact with new seasonal norms to influence ecosystems.

Here, we use a meta-ecosystem model to investigate how key seasonal characteristics—intensity of seasonal variation and duration of seasons—mediate the impact of migration on species persistence, coexistence and ecosystem function. We further investigate how responses of migratory consumers to changing seasonality can influence the ecological effects of migration. To do this, we study the model across the full suite of migration-seasonality scenarios, from migration timing and duration being perfectly matched with seasonal dynamics to strongly mismatched. Our approach explicitly integrates the temporal dynamics of ecosystems, which is key to addressing these broader ecological questions. We find that both the intensity and duration of seasonal fluctuations in primary production mediate the effects of consumer migration on species coexistence and ecosystem function, and that these outcomes depend on the timing of migration, the characteristics of competitors, and the structure of the meta-ecosystem. Importantly, phenological mismatches between migration and seasonal onset can shift coexistence patterns, modify biomass stocks at local and meta-ecosystem scales, and in certain cases, drive the extinction of migratory and non-migratory consumers. Together, these results indicate that migration and seasonality can jointly shape community structure and ecosystem functioning across scales, advancing our understanding of how time and movement interact to structure ecological systems in a changing world.

## Methods

### Meta-ecosystem models

We advanced a classical mathematical model of two coupled ecosystems (Guichard and Marleau, 2021; Marleau et al., 2010) to integrate migration and seasonality (Fig.1). Each local ecosystem comprises an inorganic nutrient compartment (*N_i_*), a primary producer compartment (*P_i_*), and depending on the model, one or more consumer compartments that are either non-migratory (*C*_1_ and *C*_2_) or migratory (*C_mig_*). Primary producers take up nutrients for growth following a Type I functional response, characterized by an uptake rate *u_p_*(*t*) as defined in (eq.3) for the seasonal ecosystem and *a*_2_ for the non-seasonal. Primary producers face losses due to mortality and other factors, with a fraction being exported from the meta-ecosystem at a rate *m_p_*. Consumption by consumers follows a Type II functional response, characterized by an attack rate *F* and handling time *h*. Consumers also experience losses through mortality at a rate *m_c_*. Non-migratory consumer compartments remain in a single ecosystem and feed on local primary producers, while the migratory consumer compartment moves between local ecosystems, consuming primary producers in the ecosystem where it is present. Both local ecosystems are open at the nutrient level, receiving a constant, independent input of nutrients *I_i_* and exporting nutrients at a rate *e_i_*. These external nutrient inputs and outputs add realism to the model, representing natural processes such as leaching, but are not crucial for the persistence of the organisms (Marleau et al., 2010). Local ecosystems exhibit identical parameter values except for the attack rates of consumer compartments, *F_i_* and *F_mig_*, which may differ between the two ecosystems according to the model’s specifications (Fig.1, also see Appendix A).

**Figure 1.**
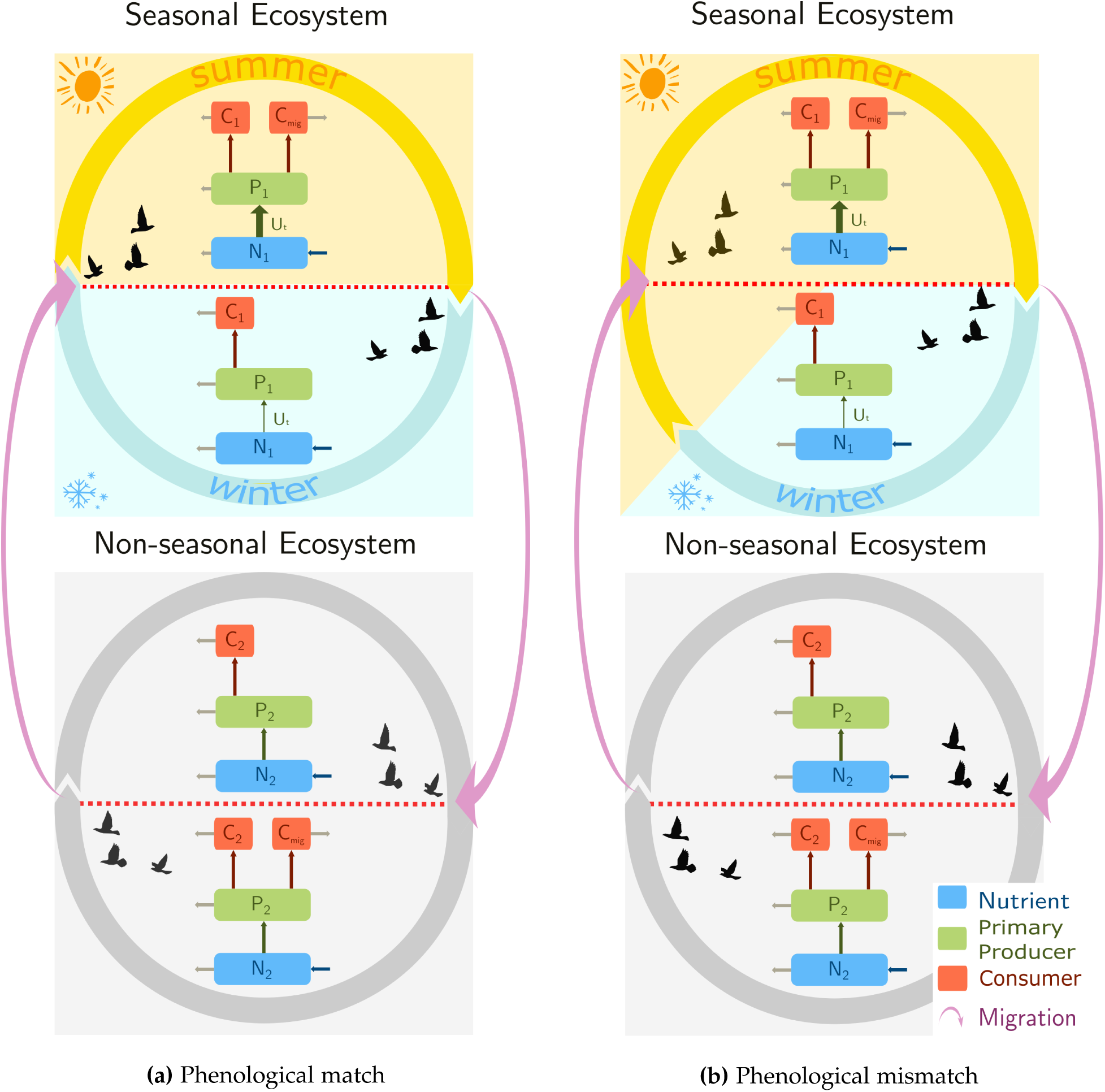
Conceptual diagram of the meta-ecosystem model with a consumer migrating between a seasonal (Ecosystem 1) and non-seasonal ecosystem (Ecosystem 2), in the case of (a) phenological match and (b) mismatch between seasonal onset and migration arrival timing. Migration is represented by bidirectional arrows: movement from the seasonal to the non-seasonal ecosystem occurs at the end of summer, while return migration from the non-seasonal to the seasonal ecosystem occurs either aligned with summer onset in the seasonal ecosystem (phenological match) or before/after summer onset (phenological mismatch). Symbols used in the figure are: *C*_1_ and *C*_2_ for the non-migrant consumers, *C_mig_*for the migrant consumer, *N_i_*for the inorganic nutrient, *P_i_*for the primary producers (*i* for each ecosystem). *U*(*t*) stands for the seasonal uptake rate of the primary production (reaching the minimum in winter and maximum in summer).

The meta-ecosystem model allows for spatial heterogeneity in seasonal dynamics. We integrate seasonal variability into Ecosystem 1, allowing seasonality to vary continuously from no seasonality to strong seasonality (Fig.1). Whereas, Ecosystem 2 is maintained at a stable equilibrium, representing no seasonal variability. We focus on this idealized case as it isolates the ecological effects of seasonality by removing any timing differences between ecosystems. Subsequent research may consider extending the present model to encompass the study of asynchronous seasonal ecosystems. For clarity, we hereafter refer to Ecosystem 1 as the seasonal ecosystem and Ecosystem 2 as the non-seasonal ecosystem. Seasonality is integrated into the seasonal ecosystem through the uptake rate of the primary producer compartment *u_p_*(*t*). Specifically, the uptake rate is modulated by a periodic function *u_p_*(*t*) (eq.3, see also Appendix B) to simulate seasonality. The characteristics of seasonality are modified by two parameters: the amplitude of oscillations *amp*_1_ which represents the intensity of the seasonal variability (Lisovski et al., 2017), and the length of summer over a year %*_summer_* which represents the period in which the uptake is above the mean value. For simplicity, we describe our analysis and findings with reference to two distinct seasons: summer and winter. From an empirical perspective, summer is intended to represent a period with higher primary production than the mean, linked to a longer photo-period and higher temperatures.

We first study the model with only the migratory consumer species (*C_mig_*) present, setting the non-migratory consumer compartments (*C*_1_ and *C*_2_) to zero. We subsequently add non-migratory consumer species to both the seasonal (*C*_1_) and non-seasonal (*C*_2_) ecosystems (Fig.1a). All consumers have identical parameter values, except for the attack rates that we vary systematically. We used a hybrid dynamical systems approach (Hutchison et al., 2024), alternatively solving the first set of differential equations where the migratory consumer is in the seasonal ecosystem (eq.1), then, solving the second set where the migratory consumer is in the non-seasonal ecosystem (eq.2). The final conditions of the previous solution become the initial conditions for the next numerical resolution. Technically, the migration between ecosystems is realized by linking the migratory consumer alternatively to each primary producer (Hutchison et al., 2024; Moisan et al., 2023). We set the migratory consumer’s departure time from the seasonal ecosystem to match the end of summer, whereas we allow the migratory consumer’s return arrival time to the seasonal ecosystem to vary in the simulations to capture mismatches in migration timing and the onset of the growing season (Fig.1b, see also Appendix c). The resulting duration of time spent in the seasonal ecosystem Δ*_mig_*, can be perfectly matched with or offset from the summer season, which enables us to introduce phenological mismatches. The dynamics of the meta-ecosystem are given by the following set of differential equations:

*Migratory consumer present in the seasonal ecosystem:*

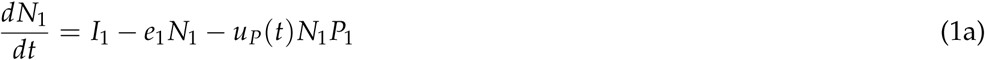

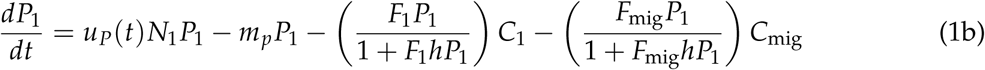

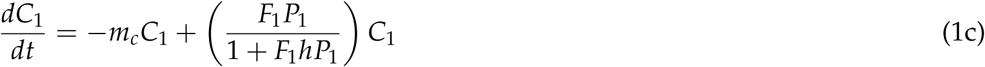

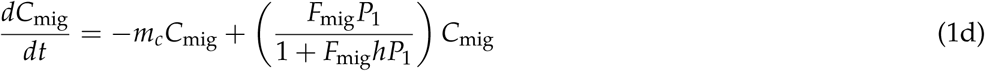

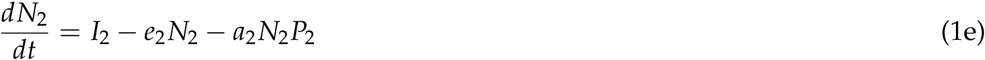

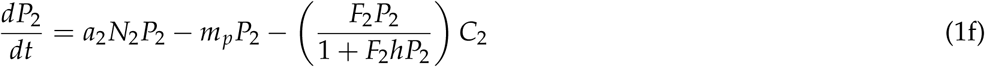

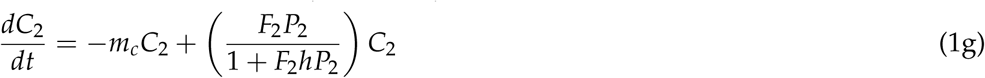

*Migratory consumer present in the non-seasonal ecosystem:*

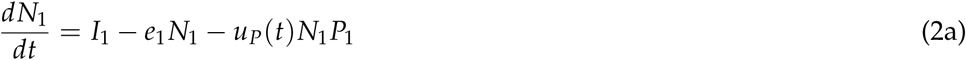

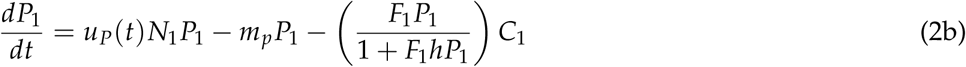

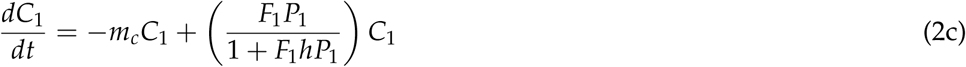

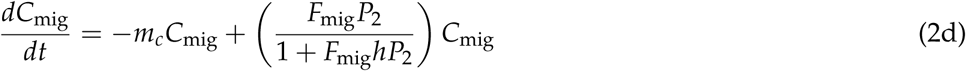

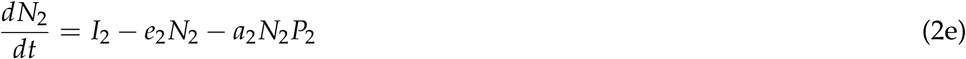

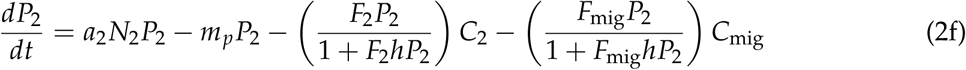

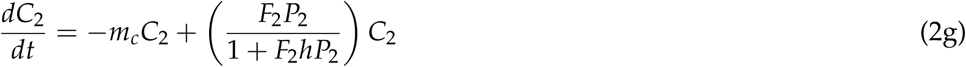

*Function modeling the seasonality of the uptake rate of primary producers in the seasonal ecosystem:*

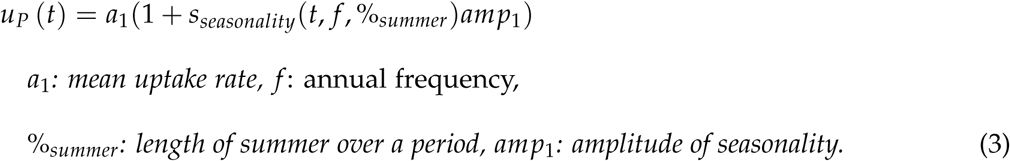

### Analysis

We first derive the non-trivial equilibria with seasonality numerically and we restrict the remainder of our analysis to parameter values that enable locally stable equilibria. We conducted sensitivity analysis of our main results according to Shaw and Levin (2011) (see Appendix D).

We conduct numerical analysis to study how migration affects consumer persistence, coexistence and ecosystem functions under a broad range in values of seasonal characteristics (*amp*_1_, %*_summer_*) and considering matches and mismatches between the start of the growing season and the timing of migration. We use an ode solver *solve ivp* (Virtanen et al., 2020) in SciPy v1.11.4 (Python 3.12.1) to illustrate the model’s dynamics, incorporating seasonality and migration, and run the simulation for 30 000 iterations. To isolate the impact of migration, we study the model with and without migration. To investigate the impact of mismatches between seasonal duration and migration timing, we independently change the length of summer and consumer arrival timing in the seasonal ecosystem keeping the departure event aligned with the end of summer (Fig.1).

#### Assessing consumers persistence and coexistence

We first study how migration affects the persistence of the migrating consumer, in the absence of non-migratory consumers in the meta-ecosystem. We define persistence as a compartment were the value of biomass is above 10^−6^, the tolerance of the numerical solver. We stepwise introduce non-migratory consumer species in the seasonal (*C*_1_) and non-seasonal ecosystem (*C*_2_) to explore how migration influences consumer coexistence. We integrate differences in competitive ability through the consumers’ attack rates, where a higher attack rate indicates greater competitive ability for resources.

#### Analyzing biomass production

We investigate the impact of migration and seasonality on the total biomass of the meta-ecosystem and the biomass of migratory consumer by summing stocks of all ecosystem compartments, or just the migrant, over a full seasonal cycle. We analyze the effect of migration on stocks across two characteristics of the migrant: competitiveness relative to non-migratory consumers (*F_mig_* − *F_i_*) and time spent in the seasonal ecosystem (Δ*_mig_*). Specifically, we calculated the relative difference in biomass with and without migration for the overall meta-ecosystem, and the absolute difference for the migrant biomass.

## Results

### Seasonal characteristics and migration timing mediate the persistence of migratory consumers

To establish a baseline, we demonstrate that migrating promotes persistence of the consumer in seasonal environments (Fig.2). Environments with relatively low amplitudes of variability in primary production allow consumer persistence both with and without migration (see Fig.2, blue zone). Whereas, in the absence of migration, high amplitudes of variability prevent consumer persistence where it can occur with migration (see Fig.2, pink zone). Only when summer makes up the majority of the year can non-migratory consumers persist when faced with high variability in primary production. Migration allows persistence where non-migratory consumers cannot persist (Fig.2a, pink zone), as migratory consumers are in the non-seasonal ecosystem during the trough of the low productivity period in the seasonal ecosystem. Introducing a phenological mismatch between seasonal primary production and the arrival date of migratory consumers hinders their persistence when seasonality is high and summers are short (Fig.2b white zone). However, the mismatch between the early arrival date and the start of summer needs to be large enough to trigger extinction.

**Figure 2.**
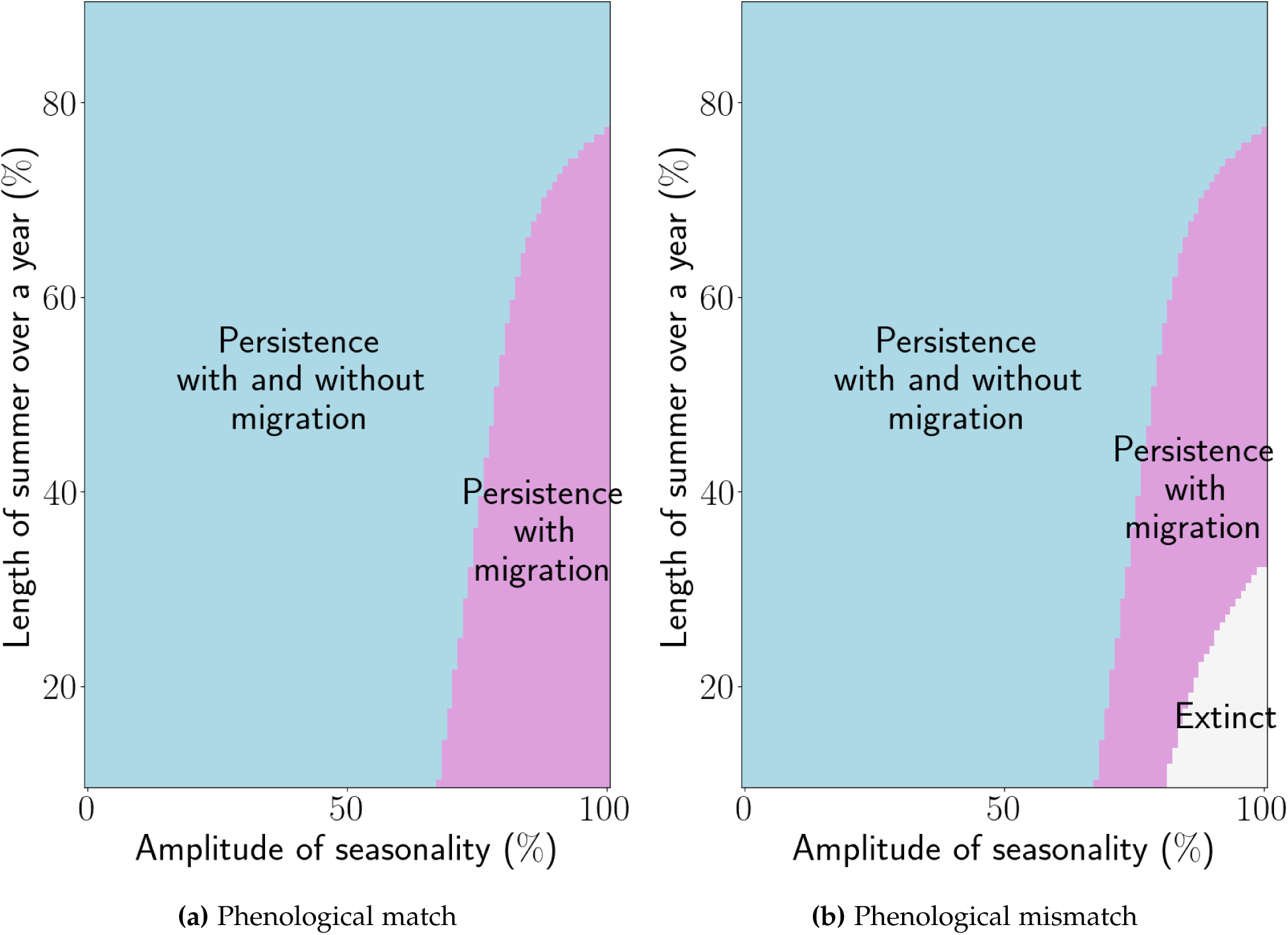
In seasonal ecosystems, the persistence of consumers can depend on migration and on either (a) phenological match or (b) mismatch between the arrival date of migratory consumers and the onset of summer. For both panels, the length of summer varies from 10% to 90% of the year. In panel (a), there is a phenological match, meaning the time spent in the seasonal ecosystem varies in accordance with the length of summer, from 10% to 90%. In panel (b), there is mismatch: the time spent in the seasonal ecosystem is fixed to 50%, while the length of summer varies. The blue zone shows the persistence of consumers with and without migration. The pink zone shows the persistence of consumers only with migration. The white zone shows where consumers cannot persist with or without migration. Parameters values used: see Appendix D.

**Figure 3.**
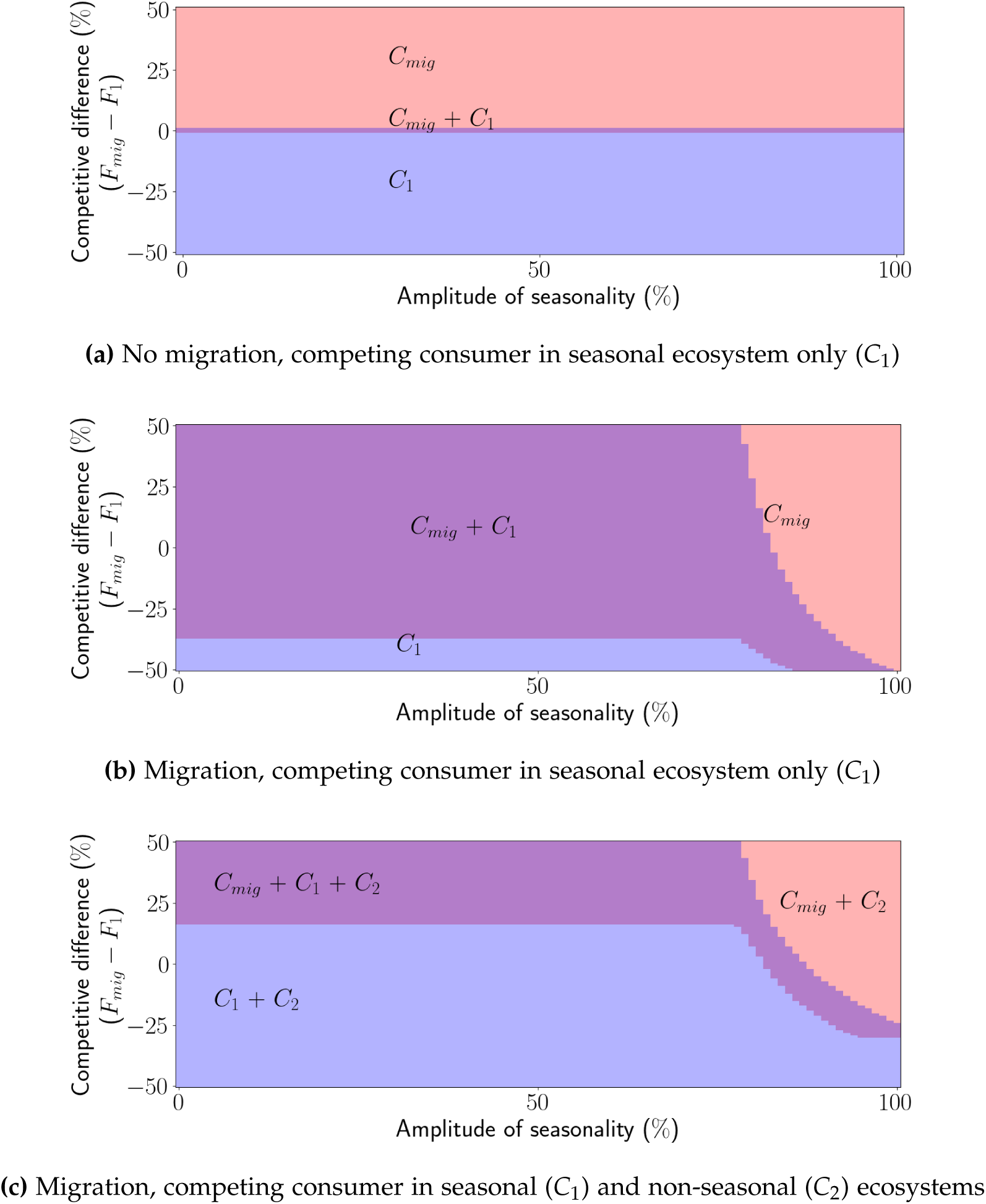
The coexistence of migratory and non-migratory consumers in a seasonal meta-ecosystem under increasing intensity (amplitude) of seasonality. Panels show the outcomes: (a) no migration, (b) migration and a non-migratory competitor (*C*_1_) in the seasonal ecosystem only, and (c) migration and a non-migratory competitor in both the seasonal (*C*_1_) and non-seasonal ecosystem (*C*_2_). Coexistence outcomes are shown across a range of competitive differences, by varying the migratory consumer’s attack rate (*F_mig_*), while keeping the attack rates of *C*_1_ and *C*_2_ constant and equivalent. A positive competitive difference indicates the migratory consumer has a higher attack rate, while a negative difference means the non-migratory consumers have the higher attack rate. In each panel, the purple zone indicates the coexistence region for *C*_1_ and *C_mig_*. The blue zone represents the region where only *C*_1_ persists, while the pink zone indicates where only *C_mig_*persists. Parameter values used: see Appendix D, length of summer = 50% of the year.

### Coexistence of consumer species within a seasonal ecosystem is influenced by migration and meta-ecosystem structure

We first present the coexistence dynamics under varying amplitudes of seasonality and differences in consumer competitive abilities, while setting the length of summer at 50% of the year and considering no mismatch between arrival timing and seasonal onset. To establish a coexistence baseline, we begin by studying the coexistence dynamics of two consumer species in the seasonal ecosystem, in the absence of migration (Fig.3a). In the absence of migration, *C*_1_ and *C_mig_* compete for resources year-round in the seasonal ecosystem and can only coexist if they have identical attack rates, effectively making them functionally identical (Fig.3a, purple zone, *F_mig_* − *F*_1_ = 0).

Next, we introduce migration into the meta-ecosystem by allowing the migratory consumer (*C_mig_*) to spend equal time in the seasonal and non-seasonal ecosystems. To gradually build up our predictions, we first include a non-migratory consumer compartment in the seasonal ecosystem only (*C*_1_), while excluding non-migratory consumers from the non-seasonal ecosystem (Fig.3b). By reducing resource use overlap in the seasonal ecosystem, migration expands the coexistence area (Fig.3b, purple zone), allowing a broader range of competitive differences between consumers. However, with higher amplitudes of variability in the seasonal ecosystem, we observe that migration can result in the exclusion of non-migratory consumers from the seasonal ecosystem, even when migratory consumers have a substantially lower attack rate than non-migratory consumers (Fig.3b, pink zone).

Finally, we add two non-migratory consumer competitors, one in the seasonal (*C*_1_) and the other in the non-seasonal (*C*_2_) ecosystem (Fig.3c), under the same assumption that the migrant spends equal time in each ecosystem. Comparing Fig.3b and Fig.3c shows how competition in the non-seasonal ecosystem modifies the coexistence area in the seasonal ecosystem. Specifically, we observe that the coexistence zone of *C*_1_ and the migratory consumer is reduced when the amplitude of seasonality relatively is low; the migratory consumer is excluded, while *C*_1_ can persist (Fig.3c, purple zone). Whereas, when the amplitude of seasonality is high, the coexistence zone of *C*_1_ and the migratory consumer is shifted toward smaller competitive differences (Fig.3c, purple zone), i.e., if the migratory consumer is a substantially weaker competitor it cannot coexist.

The presence of a consumer in the non-seasonal ecosystem similarly reduces the persistence area of the migrant (Fig.3c, pink zone), and increases the persistence area of the non-migrant in the seasonal ecosystem when the amplitude of seasonality is relatively high (*C*_1_; Fig.3c, blue zone).

### Phenological mismatches shift the coexistence of migrants with non-migratory consumers

Next, we examine how seasonal duration and phenological mismatch influences the coexistence area (Fig.4). To do so, we keep consumer attack rates constant while varying the amplitude of seasonality and the length of summer, comparing scenarios with matched (Fig.4a and 4c) and mismatched (Fig.4b and 4d) migration arrival timing. With a phenological mismatch between the migrants’ arrival timing and seasonal primary production, we observe opposing effects on the persistence of the migrant, depending on whether a non-migratory competitor is present in the non-seasonal ecosystem. Consistently, highly seasonal environments (with high amplitude and short summers) favor the migrant. When there is no competing species in the non-seasonal ecosystem (Fig.4a and 4c), we observe that a phenological mismatch does not affect the persistence of the migrant (note that for lower migrant attack rates, phenological mismatch can lead to migrant extinction for high amplitude, short summers as shown in Fig.2b). However, we observe phenological mismatch can strongly promote the persistence of the non-migratory species in the seasonal ecosystem (*C*_1_) in the case of longer summers in the seasonal ecosystem (migrants arrive late relative to onset of summer) and slightly reduce its persistence for short summers (migrants arrive early relative to onset of summer; compare purple zone in Fig.4a and in Fig.4c).

**Figure 4.**
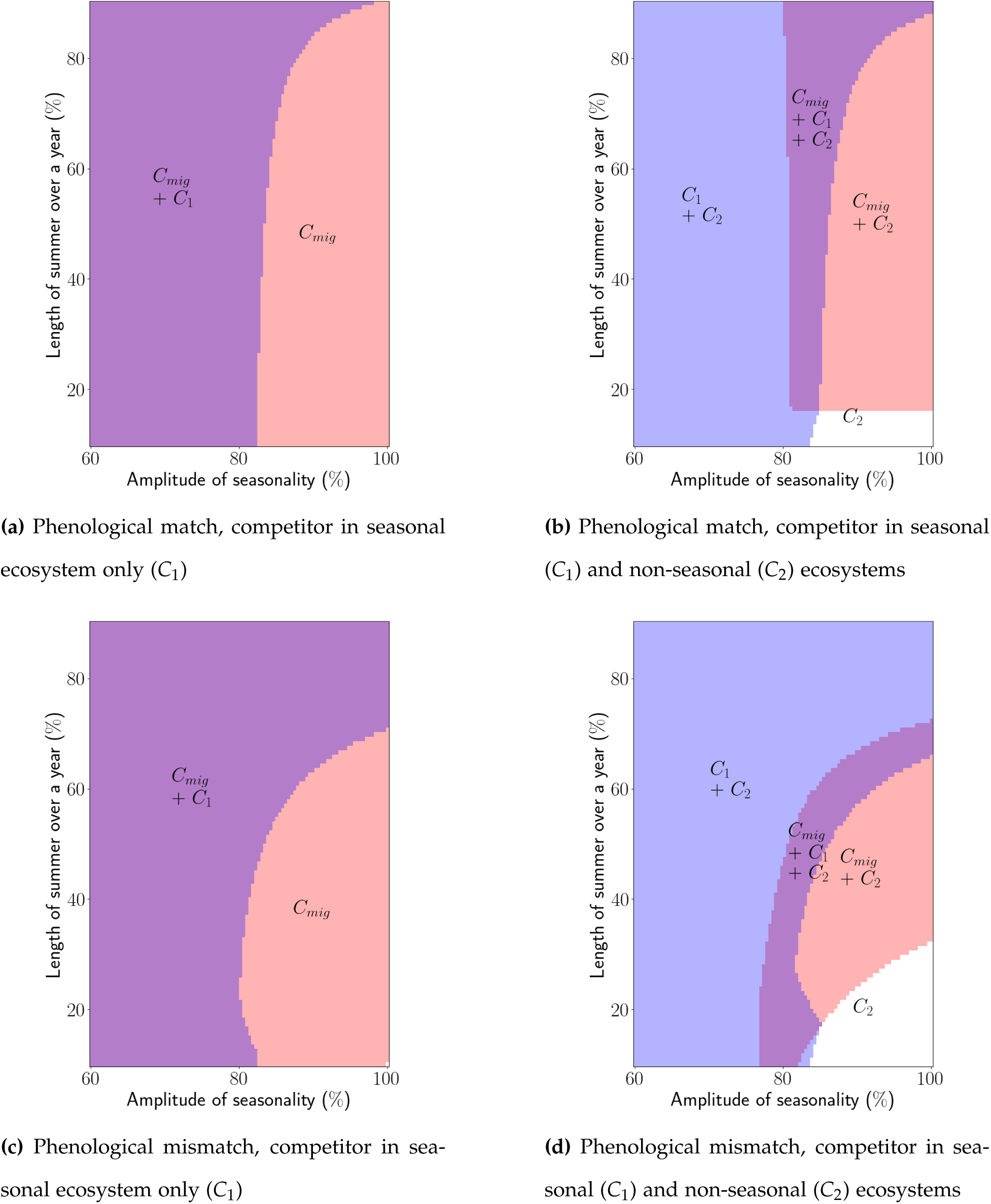
The coexistence of migratory and non-migratory consumers in a seasonal meta-ecosystem under increasing seasonal intensity (amplitude) and under (a and b) matched versus (c and d) mismatched seasonal onset and migration arrival timing (time spent in seasonal ecosystem fixed at 50%). (a and c) show coexistence outcomes with a non-migratory competitor (*C*_1_) in the seasonal ecosystem only, and (b and d) show coexistence outcomes with a non-migratory competitor in both the seasonal (*C*_1_) and non-seasonal ecosystem (*C*_2_). In each panel, the purple zone indicates the coexistence region for *C*_1_ and *C_mig_*. The blue zone represents the region where only *C*_1_ persists, while the pink zone indicates where only *C_mig_* persists. The white zone indicates where no consumer persist in the seasonal ecosystem. Parameter values used: see Appendix D.

In accordance with the coexistence results across varying consumer attack rates (Fig.3c), the presence of a non-migratory competitor in the non-seasonal ecosystem (*C*_2_) strongly modifies the seasonal parameters enabling coexistence of consumers (compare purple zone between Fig.4a with Fig.4c and Fig.4b with Fig.4d). In particular, there is a reduction in the seasonal parameters allowing the persistence of the migrant. When arrival timing is matched with seasonal onset (Fig.4b), the presence of a competitor in the non-seasonal ecosystem hinders the persistence of the migrant for lower amplitudes of seasonality. A phenological mismatch further reduces the persistence of the migrant when summer is longer (migrants arrive late relative to onset of summer)(compare pink zone Fig.4b and Fig.4d), while it promotes the persistence of the competitor in the seasonal ecosystem (*C*_1_). Conversely, when summer is shorter, phenological mismatch (migrants arrive early relative to onset of summer) can mildly promote the coexistence of the migrant with *C*_1_ and *C*_2_ when the amplitude of seasonality is at intermediate levels (compare purple zone Fig.4b and Fig.4d) and reduce the persistence of the migrant when seasonality is high (compare white zones).

### Consumer competitiveness and meta-ecosystem structure mediate how migration timing affects biomass production

To investigate the effects of migration on biomass distribution at both meta-ecosystem and local scales, we varied key characteristics of migrants: their attack rate and their arrival date to the seasonal ecosystem, the latter of which we did by varying the proportion of time migrants spent in the seasonal ecosystem throughout the year while keeping seasonal duration constant. For this fixed seasonal baseline (with an amplitude of seasonality of 80% , a length of summer adding up to 50% of the year, and no local competitors in either ecosystem), we compared biomass with and without migration for the entire meta-ecosystem, and for migrant consumers, alone. We found that migration has the strongest positive effect on the biomass of the migrant when the attack rate of the migrant is lower than the non-migratory competitors (Fig.5a dark orange zone), which also generally corresponds to a positive increase in biomass of the meta-ecosystem (Fig.5b, light orange zone). However, when the migratory consumer has a higher attack rate than its non-migratory competitors, migration positively impacts the biomass of the migratory consumer but generally does not result in increased biomass at the meta-ecosystem scale (Fig.5, purple zone).

**Figure 5.**
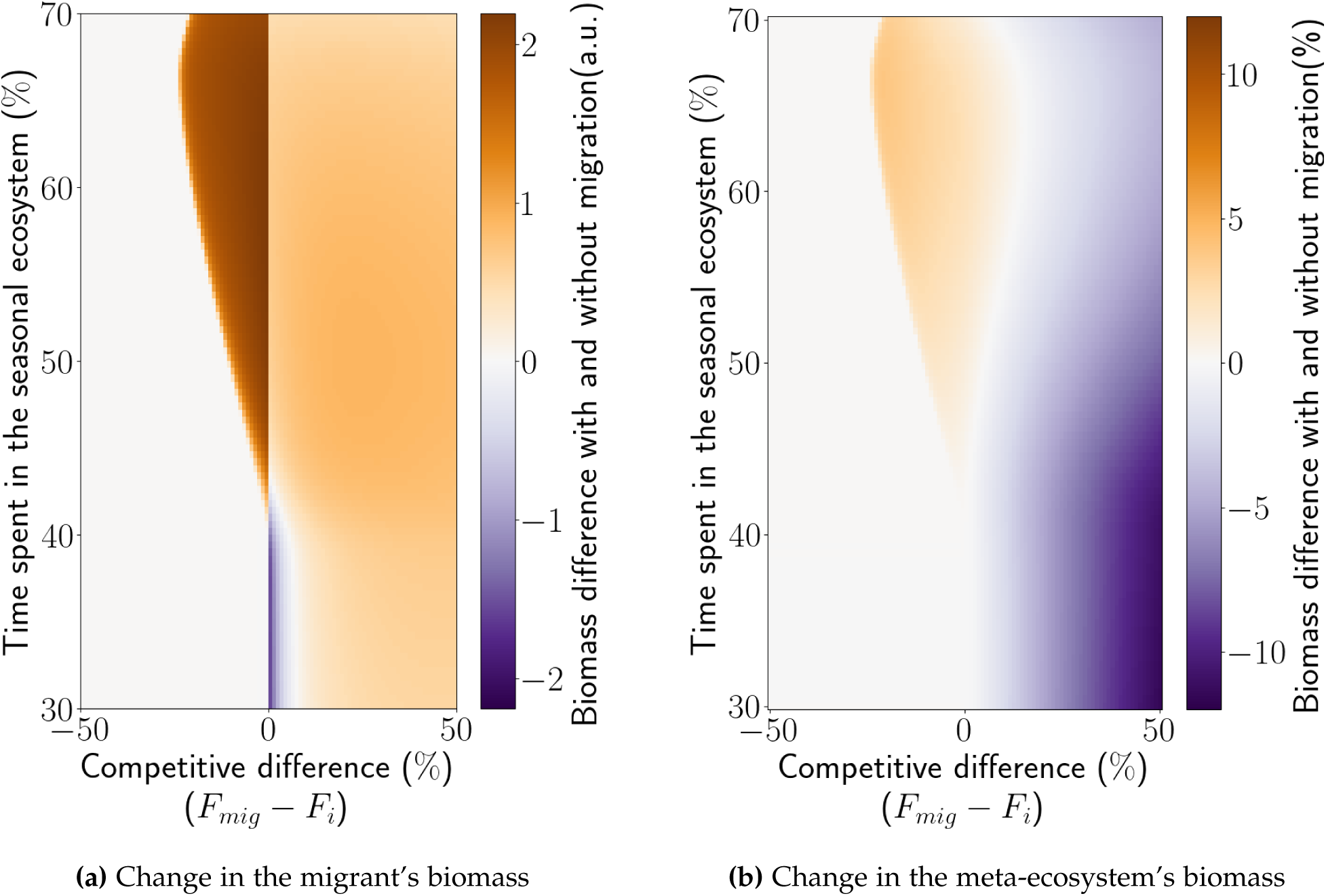
The stocks of biomass of (a) the migratory consumer and (b) the entire meta-ecosystem, are influenced by the migrant’s characteristics: time spent in each ecosystem and attack rate. In each panel, orange indicates a positive effect of migration, while purple shows a negative effect. Both panels show the result when non-migratory consumers are present in both the seasonal and non-seasonal ecosystems. (a) shows the biomass difference for the migrant consumer, with and without migration. (b) shows the percentage difference in biomass for the entire meta-ecosystem, with and without migration. Parameter values used: see Appendix D, amplitude of seasonality = 80% and length of summer = 50% of the year.

We found this effect of migration on biomass in the meta-ecosystem and in the stocks of the migratory consumer to also be impacted by the arrival date to the seasonal ecosystem (time spent in the seasonal ecosystem) (Fig.5 and Fig.6). Specifically, early arrival dates relative to seasonal onset (time spent in seasonal ecosystem is more than 50 %) expand the potential for migration to have positive impacts on biomass (or reduce negative impacts). Whereas, late arrival dates relative to seasonal onset (time spent in seasonal ecosystem is less than 50 %) can eliminate the positive effects of migration when migrants have lower attack rates than non-migrants. When migrants have higher attack rates than non-migrants, late arrival dates generally lead to negative impacts on biomass: lower consumer biomass and meta-ecosystem biomass with migration.

**Figure 6.**
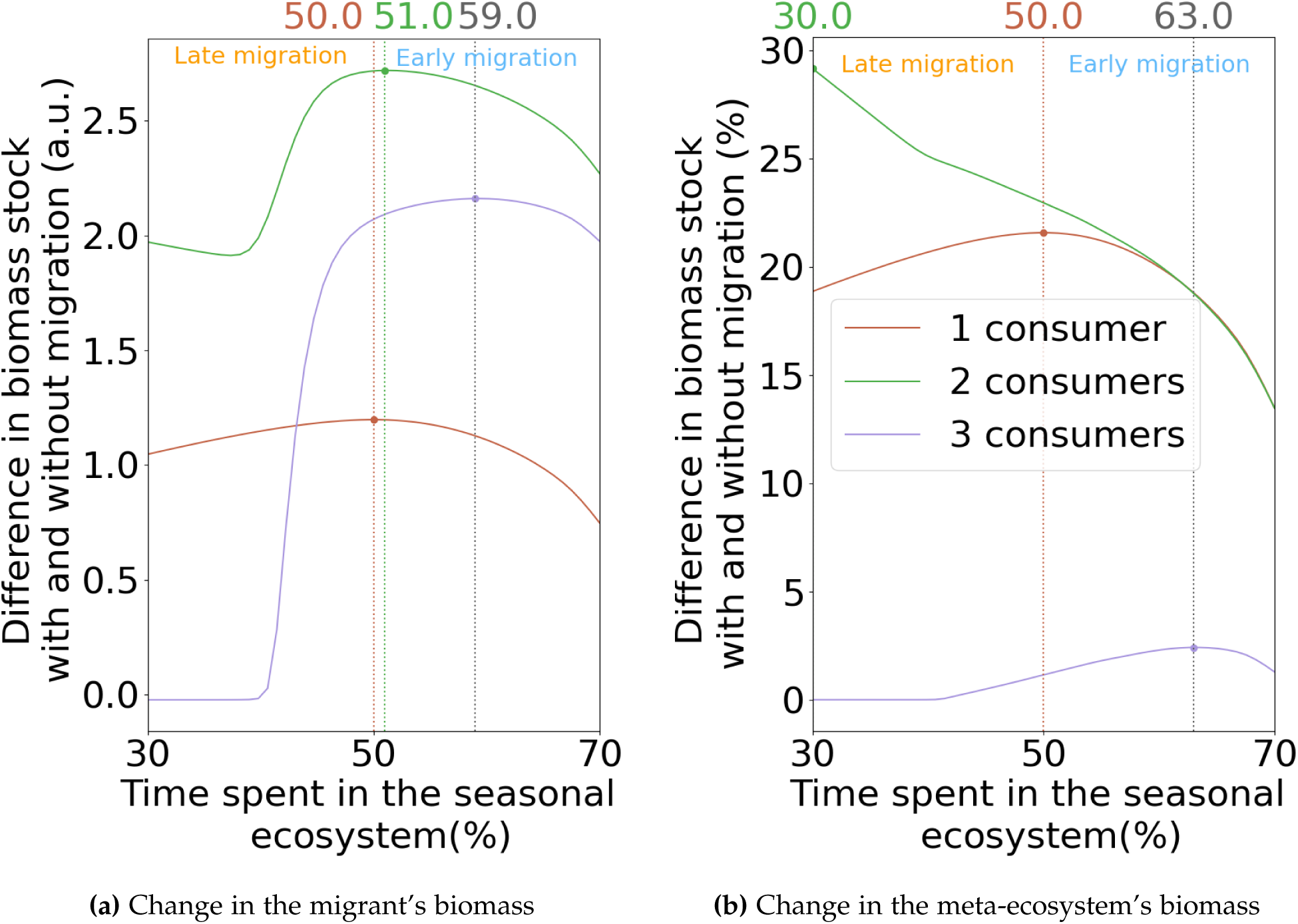
The stocks of biomass with versus without migration at (a) the migratory consumer and (b) the entire meta-ecosystem, with the migratory consumer being the only consumer species in the meta-ecosystem (brown), with the migratory consumer and a competitor (*C*_1_) in the seasonal ecosystem only (green), and with the migratory consumer and competitors in both the seasonal (*C*_1_) and non-seasonal (*C*_2_) ecosystems (purple). Both panels show the outcome for a single competitive difference between the migratory and non-migratory consumers (*F_mig_* − *F_i_* = 0%). The optimal arrival timing (in terms of biomass gain) of the migrant depends on competition with the non-migrant consumers. Parameter values used: see Appendix D, amplitude of seasonality = 80% and length of summer = 50% of the year.

Notably, we found that the arrival date maximizing the biomass for both the migrant and the overall meta-ecosystem stocks depends on the presence of non-migratory competitors (Fig.6). In the absence of non-migratory consumers in both ecosystems, the arrival date which has the strongest positive impact on biomass of the migratory consumer and entire meta-ecosystem is synchronized with the start of summer (Fig.6b, brown line). When a non-migratory consumer species is present in the seasonal ecosystem only, the arrival date which has the strongest positive impact at the migrant level (Fig.6a, light green line) shifts slightly to before the start of summer. However, at the meta-ecosystem level migration has the most positive effect the later migrants arrive to the seasonal ecosystem (Fig.6b, light green line). Whereas, when non-migratory consumer species are present in both the seasonal and non-seasonal ecosystems, the arrival date which has the strongest positive impact on the migrant and the meta-ecosystem (Fig.6, light purple line) strongly shifts to before the start of summer in the seasonal ecosystem.

## Discussion

We studied how consumer migration interacts with seasonality to influence consumer coexistence and ecosystem functions within a meta-ecosystem. Our findings demonstrate that migration can promote the persistence of less competitive consumers in seasonal ecosystems, both migratory and non-migratory, thus fostering species coexistence. However, we found the impacts of migration depend critically on the characteristics of seasonality and the timing of migration relative to seasonal dynamics; phenological mismatches between migrant arrival and the start of summer were capable of mediating the effects of migration on migrant persistence, consumer coexistence, and on meta-ecosystem biomass. Further, we found that species interactions in one ecosystem visited by migrants can mediate the influence of arrival times and coexistence outcomes in another ecosystem, underscoring the significance of considering meta-ecosystem dynamics in migration studies. Understanding how the temporal characteristics of ecological processes, as well as phenological mismatches (Post and Forchhammer, 2007; Rudolf, 2019; Visser and Gienapp, 2019), can modify ecological dynamics and functions has been emphasized as a key challenge in ecology (Hastings, 2010; Hernández-Carrasco et al., 2025; Yin and Rudolf, 2024). Moreover, while migration is a widespread and well-studied phenomenon at the population level, we still largely lack a mechanistic understanding of how it influences biodiversity and ecosystem functions across scales (Bauer and Hoye, 2014; Peller et al., 2023). Our study provides fundamental theoretical insight into how migration shapes the dynamics of interconnected ecosystems, offering a mechanistic foundation for understanding its complex role amid shifting seasonal patterns.

Our model predicts that characteristics of seasonality—its amplitude and duration—can influence migrant persistence and the coexistence of migrant and non-migrant consumer species in meta-ecosystems. Migration promotes the persistence of consumers inhabiting strongly seasonal ecosystems, agreeing with empirical observations (Acker et al., 2021; Hurlbert and Haskell, 2003; Milner-Gulland et al., 2011). In particular, migration enables the consumer population to never face the most extreme resource conditions that occur in highly seasonal ecosystems (Hutchison et al., 2024). Notably, our model further predicts migration can enable migrants to exclude non-migratory competitors in strongly seasonal ecosystems despite being competitively inferior for local resources. Longer summers, however, interact with seasonal intensity, generally making migration a less competitive strategy. In fact empirical results show that longer summer lead to a smaller proportion of migrants (Martin et al., 2018). This is explained in our model because longer summers benefit resident, non-migratory consumers, allowing them to increase in density and out-compete migratory consumers (Payo-Payo et al., 2022; Van Moorter et al., 2020). Interactions between migrants and residents are increasingly recognized as an important driver of community dynamics, yet their mechanisms remain poorly understood, particularly regarding the impacts of migrants on residents and the mediating role of seasonality (Navarro-Velez and Dhondt, 2025). By systematically varying key seasonal parameters (Lisovski et al., 2017; Torstenson and Shaw, 2025), our study provides mechanistic insights into how natural variation and changes in seasonality can alter the ecological impacts of migration and reshape communities across meta-ecosystems.

We show that mismatches between seasonal onset and migration timing can mediate the effects of migration on the coexistence of migratory and non-migratory consumers and meta-ecosystem biomass. This suggests that the consequences of altered migration timing extend beyond population-level outcomes to affect meta-ecosystem structure and function, advancing theoretical predictions of cross-ecosystem coupling (Gounand et al., 2018; Leroux and Loreau, 2012; Loreau et al., 2003). Importantly, the consequences of these mismatches depend on the structure of the meta-ecosystem. In simpler meta-ecosystems, where the non-seasonal ecosystem lacks a competing consumer species, mismatches in migration timing can alleviate competition for both consumers in the seasonal ecosystem through niche partitioning in time and space (Salewski et al., 2003). Arriving late to the seasonal ecosystem promotes coexistence by reducing direct competition in the seasonal ecosystem during periods of peak resource availability. However, with increasing food web complexity—i.e., via the addition of a competing consumer species in the non-seasonal ecosystem—optimal synchronization between migration timing and seasonal primary production becomes crucial for migrant persistence and consumer coexistence. Migrants, in particular, directly face multiple competitors and must time their exploitation of resource peaks with precision to remain competitive. Arriving late can prevent the migrant from coexisting with the non-migrants by being out-competed in terms of density, a result that aligns with empirical evidence of trophic mismatches in plant–insect–bird systems under climate change (Kharouba et al., 2018; Visser and Both, 2005). Phenological mismatches have been observed to have variable outcomes (Post and Forchhammer, 2007) and have primarily been studied in the context of single pairwise interactions (Beard et al., 2019; Visser and Gienapp, 2019). By demonstrating species interactions in the different ecosystems migrants connect can shape the outcomes of phenological mismatches, our findings support the need for a broader meta-ecosystem perspective on phenological change (Rudolf, 2019).

Our study predicts that migration-driven coexistence does not always lead to enhanced biomass at the meta-ecosystem level, aligning with empirical findings (Moisan et al., 2023). For instance, when migrants arrive at the onset of summer and spend equal amounts of time in both ecosystems, migration promotes consumer diversity. However, this increase in diversity can result in lower total biomass across the meta-ecosystem if the migrant is the superior competitor. In this case, greater consumer diversity diminishes the efficiency of primary production consumption, cascading into an overall decrease in consumer biomass across ecosystems while simultaneously increasing primary producer biomass. Theoretical studies, such as Marleau and Guichard (2019), align with this observation, suggesting that movements of organisms can promote coexistence while altering ecological functions. Similarly, Brodersen et al. (2008) showed that migration can exert opposing top-down effects on primary production depending on the timing and density of migrants. In our study, we found that for migration to positively impact ecosystem biomass, the migrant generally needs to have a lower attack rate than the non-migrant consumers. This allows non-migrant consumers to use primary production more consistently, while the migrant capitalizes on the seasonal abundance of resources. With a larger migratory population arriving in the seasonal ecosystem, migrants can efficiently exploit seasonal primary production without overshadowing the non-migrant consumers’ access to resources, as the latter have a higher attack rate. These findings underscore the dual impact of migration in shaping both biodiversity and ecosystem functioning, and they highlight the importance of consumer traits in mediating the outcomes (Leroux and Schmitz, 2025; Peller and Altermatt, 2024).

While it is well established that ”timing is crucial” (Yin and Rudolf, 2024), the ecological mechanisms that determine optimal arrival timing—particularly in the context of biomass production across migratory systems—remain poorly understood (Beard et al., 2019; Lameris et al., 2018; Mayor et al., 2017). We found that competition for seasonal food resources between migrants and non-migrants is a key factor shaping the optimal timing of migration with regards to maximizing migrant and meta-ecosystem biomass. In a simple scenario without competing non-migratory consumer species, the optimal arrival date for migrants to the seasonal ecosystem aligns with the start of summer to maximize biomass. However, when non-migratory species competing for the same resources are present in each ecosystem, the optimal arrival date shifts earlier. This difference occurs because migrants that avoided winter in the seasonal ecosystem have higher densities upon returning to the seasonal ecosystem than non-migrants—a carryover effect from wintering in the non-seasonal ecosystem. This numeric advantage allows migrants to better exploit seasonal resources through a slight timing offset, despite having a lower per capita attack rate. Consequently, competition for resources can favor early migration timing relative to the phenology of primary production, supporting previous theoretical predictions (Morrison et al., 2019). While previous studies have emphasized the breeding advantages of early arrival, our findings indicate that local competition for food is also crucial for understanding temporal timing in ecology (Shaw, 2016).

Our study unraveled a ”Pandora’s box” of dynamics in a time-dependent, non-equilibrium meta-ecosystem model. By introducing a periodic process in the form of growth in uptake, we demonstrated how time-varying ecological processes, can alter community dynamics (Holt, 2008; Leroux and Loreau, 2012; McMeans et al., 2015; White and Hastings, 2020) and permit coexistence that would otherwise be unattainable. In this context, we studied the dynamics of out-of-equilibrium communities, where migration stabilizes the seasonal ecosystem while destabilizing the non-seasonal one. Our study provides a critical foundation integrating migration and seasonal dynamics into metacommunity and meta-ecosystem theory (see also Hutchison et al. (2020); Moisan et al. (2025)), laying the ground work for crucial avenues for further research. As a first step to understand the impacts of migration and seasonality, we did not account for nutrient transport or recycling. Migrants, however, are known to be transporters of nutrients between ecosystems (Bauer and Hoye, 2014; Brandt et al., 2024). Including nutrient recycling could reveal cascading effects of migration on nutrient stocks and primary production within both seasonal and non-seasonal ecosystems (Peller et al., 2023). These effects could emerge from resource inputs associated with nonliving migrant biomass, potentially creating pulses of nutrients that ripple through the system.

## Conclusion

We integrated seasonality and migration—into a meta-ecosystem model to examine how the interaction between these ubiquitous features of nature alter ecosystems and influence their dynamics and functioning. Although seasonality and migration are changing around the world (Hernández-Carrasco et al., 2025; UNEP-WCMC, 2024), there is still limited knowledge on how they shape entire communities and ecosystems across scales. Our study highlighted the pivotal role of migration in the coexistence of consumer species and the overall biomass of a meta-ecosystem. However, it further revealed the strong role of seasonal characteristics in mediating the impacts of migration and its timing. Further research in this direction is vital for understanding how biodiversity and ecosystem processes are being shaped by changing seasonal and migratory dynamics.

## Acknowledgments

Funding is from the Swiss National Science Foundation (grant 310030 197410) to F.A.

## 1 Statement of Authorship

All authors designed the study. LC performed the simulations and analysed the data. LC and TP wrote the first draft of the manuscript. All authors contributed to revisions.

## 2 Data Availability Statement

No data were used in this article. The code is available in the Zenodo repository: https://doi.org/10.5281/zenod

## Literature Cited

P. Acker, F. Daunt, S. Wanless, S.J. Burthe, M.A. Newell, M.P. Harris, H. Grist, J. Sturgeon, R.L. Swann, C. Gunn, A. Payo-Payo, and J.M. Reid. Strong survival selection on seasonal migration versus residence induced by extreme climatic events. Journal of Animal Ecology, 90(4):796–808, 2021.

F. Bairlein. Migratory birds under threat. Science, 354:547–548, 11 2016.

K. Bandara, O. Varpe, L. Wijewardene, V. Tverberg, and K. Eiane. Two hundred years of zooplankton vertical migration research. Biological Reviews, 96(4):1547–1589, 2021.

S. Bauer and B.J. Hoye. Migratory animals couple biodiversity and ecosystem functioning world-wide. Science, 344:1242552, 4 2014.

K.H. Beard, K.C. Kelsey, A.J. Leffler, and J.M. Welker. The missing angle: Ecosystem consequences of phenological mismatch. Trends in Ecology and Evolution, 34:455–457, 6 2019.

P Berthold. Control of Bird Migration. Springer Science and Business Media, 1996.

C. Both and M.E. Visser. Adjustment to climate change is constrained by arrival date in a long-distance migrant bird. Nature, 411:296–298, 5 2001.

M. Brandt, D. Gominski, F. Reiner, A. Kariryaa, V. B. Guthula, P. Ciais, X. Tong, W. Zhang, D. Ortiz-Gonzalo, and R. Fensholt. Severe decline in large farmland trees in india over the past decade. Nature Sustainability, 7(7):860–868, 5 2024.

J. Brodersen, E. Ådahl, C. Brönmark, and L.-A. Hansson. Ecosystem effects of partial fish migration in lakes. Oikos, 117:40–46, 1 2008.

K.K. Clausen and P. Clausen. Earlier arctic springs cause phenological mismatch in long-distance migrants. Oecologia, 173:1101–1112, 11 2013.

J. Cohen, L. Agel, M. Barlow, C.I. Garfinkel, and I White. Linking arctic variability and change with extreme winter weather in the united states. Science, 373:1116–1121, 9 2021.

J. M. Cohen, M. J. Lajeunesse, and J. R. Rohr. A global synthesis of animal phenological responses to climate change. Nature Climate Change, 8:224–228, 3 2018.

D.O. Conover. Seasonality and the scheduling of life history at different latitudes. Journal of Fish Biology, 41:161–178, 1992.

M. Doiron, P. Legagneux, G. Gauthier, and E. Lévesque. Trophic mismatch and ecosystem functioning: Can late snowmelt lead to a mismatch between predators and resources? Global Change Biology, 21(8):2572–2584, 2015.

J. Doležal, V. Lanta, O. Mudrák, and J. Lepš. Seasonality promotes grassland diversity: Interactions with mowing, fertilization and removal of dominant species. Journal of Ecology, 107: 203–215, 1 2019.

J.G. Ernakovich, K.A. Hopping, A.B. Berdanier, R.T. Simpson, E.J. Kachergis, H. Steltzer, and M.D. Wallenstein. Predicted responses of arctic and alpine ecosystems to altered seasonality under climate change. Global Change Biology, 20:3256–3269, 10 2014.

I. Gounand, C. J. Little, E. Harvey, and F. Altermatt. Cross-ecosystem carbon flows connecting ecosystems worldwide. Nature Communications, 9(1):4825, 2018.

F. Guichard and J. Marleau. Meta-Ecosystems. Springer International Publishing, 2021.

B. Haest, O. Hüppop, and F. Bairlein. Weather at the winter and stopover areas determines spring migration onset, progress, and advancements in afro-palearctic migrant birds. Proceedings of the National Academy of Sciences of the United States of America, 117:17056–17062, 7 2020.

J.H. Hansen, C. Skov, H. Baktoft, N. Jepsen, A. Koed, and J. Brodersen. Ecological consequences of animal migration: Prey partial migration affects predator ecology and prey communities. Ecosystems, 23:292–306, 2020.

G. Harris, S. Thirgood, J.G.C. Hopcraft, J.P.G.M. Cromsigt, and J. Berger. Global decline in aggregated migrations of large terrestrial mammals. Endangered Species Research, 7:55–76, 4 2009.

A. Hastings. Timescales, dynamics, and ecological understanding. Ecology, 91(12):3471–3480, 2010.

D. Hernández-Carrasco, J.M. Tylianakis, D.A. Lytle, and J.D. Tonkin. Ecological and evolutionary consequences of changing seasonality. Science, 388(6750), 2025.

R.D. Holt. Theoretical perspectives on resource pulses. Ecology, 89:671–681, 3 2008.

A. H. Hurlbert and J. P. Haskell. The effect of energy and seasonality on avian species richness and community composition. The American Naturalist, 161(1):83–97, 2003.

C. Hutchison, F. Guichard, P. Legagneux, G. Gauthier, J. Bêty, D. Berteaux, D. Fauteux, and D. Gravel. Seasonal food webs with migrations: multi-season models reveal indirect species interactions in the canadian arctic tundra. Philosophical Transactions of the Royal Society A, 378: 20190354, 8 2020.

C. Hutchison, D. Gravel, and F. Guichard. A hybrid dynamical system approach to predicting the resilience of community dynamics with seasonal migrations under climate change. Theoretical Ecology, 17:143–154, 2024.

H. M. Kharouba, J. Ehrlén, A. Gelman, K. Bolmgren, J. M. Allen, S. E. Travers, and E. M. Wolkovich. Global shifts in the phenological synchrony of species interactions over recent decades. Proceedings of the National Academy of Sciences, 115(20):5211–5216, 2018.

T.K. Lameris, H.P. van der Jeugd, G. Eichhorn, A.M. Dokter, W. Bouten, M.P. Boom, K.E. Litvin, B.J. Ens, and B.A. Nolet. Arctic geese tune migration to a warming climate but still suffer from a phenological mismatch. Current Biology, 28:2467–2473.e4, 8 2018.

S.J. Leroux and M. Loreau. Dynamics of reciprocal pulsed subsidies in local and meta-ecosystems. Ecosystems, 15:48–59, 1 2012.

S.J. Leroux and O.J. Schmitz. Integrating network and meta-ecosystem models for developing a zoogeochemical theory. Ecology Letters, 28:1–15, 2 2025.

W. Lin and C. Wang. Longer summers in the northern hemisphere under global warming. Climate Dynamics, 58:2293–2307, 5 2022.

S. Lisovski, M. Ramenofsky, and J. C. Wingfield. Defining the degree of seasonality and its significance for future research. Integrative and Comparative Biology, 57:934–942, 11 2017.

M. Loreau, N. Mouquet, and R.D. Holt. Meta-ecosystems: a theoretical framework for a spatial ecosystem ecology. Ecology Letters, 6(8):673–679, 2003.

J. N. Marleau, M. Hébert, N. Rooney, L. Xu, and K. S. McCann. Nutrient flows between ecosystems can destabilize simple meta-ecosystem modules. Journal of Theoretical Biology, 267:727–739, 11 2010.

J.N. Marleau and F. Guichard. Meta-ecosystem processes alter ecosystem function and can promote herbivore-mediated coexistence. Ecology, 100:e02699, 6 2019.

J. Martin, V. Tolon, N. Morellet, H. Santin-Janin, A. Licoppe, C. Fischer, J. Bombois, P. Patthey, E. Pesenti, D. Chenesseau, and S. Said. Common drivers of seasonal movements on the migration–residency behavior continuum in a large herbivore. Scientific Reports, 8(1):7631, 5 2018.

S.J. Mayor, R.P. Guralnick, M.W. Tingley, J. Otegui, J.C. Withey, S.C. Elmendorf, M.E. Andrew, S. Leyk, I.S. Pearse, and D.C. Schneider. Increasing phenological asynchrony between spring green-up and arrival of migratory birds. Scientific Reports, 7:1–10, 5 2017.

B. C. McMeans, K.S. McCann, M. Humphries, N. Rooney, and A.T. Fisk. Food web structure in temporally-forced ecosystems. Trends in Ecology and Evolution, 30:662–672, 11 2015.

E.J. Milner-Gulland, J.M. Fryxell, and A.R.E. Sinclair. Animal Migration: A Synthesis. Oxford University Press, 01 2011.

L. Moisan, D. Gravel, P. Legagneux, G. Gauthier, D.J. Léandri-Breton, M. Somveille, J.F. Therrien, J.F. Lamarre, and J. Bêty. Scaling migrations to communities: An empirical case of migration network in the arctic. Frontiers in Ecology and Evolution, 10:1077260, 1 2023.

L. Moisan, D. Gravel, G. Gauthier, P. Legagneux, and J. Bety. Arctic migrations shape global meta-communities: Contrasting insights from species occurrence, abundance and biomass. Global Ecology and Biogeography, 34(6):e70074, 2025.

C. A. Morrison, J.A. Alves, T.G. Gunnarsson, B. Thórisson, and J.A. Gill. Why do earlier-arriving migratory birds have better breeding success? Ecology and Evolution, 9:8856–8864, 8 2019.

K. C. Navarro-Velez and A. A. Dhondt. Wintering together: Do migrants impact residents? a literature review. Ecology and Evolution, 15(2):e70868, 2025.

Ian Newton. Chapter 1 - Introduction. Academic Press, 2007.

A. Payo-Payo, F. Pulido, V. Renaud, P. Berthier, P. Rodriguez, J. Lemaitre, and F. Cagnacci. Modelling the responses of partially migratory metapopulations to changing seasonal migration rates. Journal of Animal Ecology, 91(5):1249–1262, 2022.

T. Peller and F. Altermatt. Invasive species drive cross-ecosystem effects worldwide. Nature Ecology and Evolution, 8(6):1087–1097, 2024.

T. Peller, F. Guichard, and F. Altermatt. The significance of partial migration for food web and ecosystem dynamics. Ecology Letters, 26:3–22, 1 2023.

E. Post and M.C. Forchhammer. Climate change reduces reproductive success of an arctic herbivore through trophic mismatch. Philosophical Transactions of the Royal Society B: Biological Sciences, 363:2369–2373, 11 2007.

V.H.W. Rudolf. The role of seasonal timing and phenological shifts for species coexistence. Ecology Letters, 22:1324–1338, 8 2019.

V. Salewski, F. Bairlein, and B. Leisler. Niche partitioning of two palearctic passerine migrants with afrotropical residents in their west african winter quarters. Behavioral Ecology, 14(4):493– 502, 2003.

A.K. Shaw. Drivers of animal migration and implications in changing environments. Evolutionary Ecology, 30:991–1007, 12 2016.

A.K. Shaw and S.A. Levin. To breed or not to breed: a model of partial migration. Oikos, 120: 1871–1879, 12 2011.

A. L. Subalusky, C. L. Dutton, E. J. Rosi, and D. M. Post. Annual mass drownings of the serengeti wildebeest migration influence nutrient cycling and storage in the mara river. Proceedings of the National Academy of Sciences, 114(29):7647–7652, 2017.

M.S. Torstenson and A.K. Shaw. Strength of seasonality and type of migratory cue determine the fitness consequences of changing phenology for migratory animals. Oikos, 134:e10862, 1 2025.

UNEP-WCMC. State of the world’s migratory species report 2024. Technical report, UNEP-WCMC, 2024.

B. Van Moorter, S. Engen, J.M. Fryxell, M. Panzacchi, E.B. Nilsen, and A. Mysterud. Consequences of barriers and changing seasonality on population dynamics and harvest of migratory ungulates. Theoretical Ecology, 13(4):595–605, 2020.

P. Virtanen, R. Gommers, T.E. Oliphant, M. Haberland, T. Reddy, D. Cournapeau, E. Burovski, P. Peterson, W. Weckesser, J. Bright, S.J. van der Walt, M. Brett, J. Wilson, K.J. Millman, N. Mayorov, A.R.J. Nelson, E. Jones, R. Kern, E. Larson, C.J. Carey, İ. Polat, Y. Feng, E.W. Moore, J. VanderPlas, D. Laxalde, J. Perktold, R. Cimrman, I. Henriksen, E.A. Quintero, C.R. Harris, A.M. Archibald, A.H. Ribeiro, F. Pedregosa, P. van Mulbregt, and SciPy 1.0 Contributors. SciPy 1.0: Fundamental Algorithms for Scientific Computing in Python. Nature Methods, 17:261–272, 2020.

M. E. Visser and C. Both. Shifts in phenology due to global climate change: the need for a yardstick. Proceedings of the Royal Society B: Biological Sciences, 272(1581):2561–2569, 2005.

M. E. Visser and P. Gienapp. Evolutionary and demographic consequences of phenological mismatches. Nature Ecology and Evolution, 3:879–885, 2019.

G. Walther, E. Post, P. Convey, A. Menzel, C. Parmesan, T. J. C. Beebee, J. Fromentin, O. Hoegh-Guldberg, and F. Bairlein. Ecological responses to recent climate change. Nature, 416:389–395, 2002.

D. H. Ward, J. Helmericks, J. W. Hupp, L. Mcmanus, M. Budde, D. C. Douglas, and K. D. Tape. Multi-decadal trends in spring arrival of avian migrants to the central arctic coast of alaska: effects of environmental and ecological factors. Journal of Avian Biology, 47:197–207, 3 2016.

E. R. White and A. Hastings. Seasonality in ecology: Progress and prospects in theory. Ecological Modelling, 44:100867, 2020.

H. Yin and V.H.W. Rudolf. Time is of the essence: A general framework for uncovering temporal structures of communities. Ecology Letters, 27(7):e14481, 2024.

